# Analysis of microvascular thrombus mechanobiology with a novel particle-based model

**DOI:** 10.1101/2021.06.07.447380

**Authors:** Anastasia A. Masalceva, Valeriia N. Kaneva, Mikhail A. Panteleev, Fazoil Ataullahanov, Vitaly Volpert, Ilya Afanasyev, Dmitry Yu. Nechipurenko

## Abstract

Platelet accumulation at the site of vascular injury is regulated by soluble platelet agonists, which induce various types of platelet responses, including integrin activation and granule secretion. The interplay between local biochemical cues, mechanical interactions between platelets and macroscopic thrombus dynamics is poorly understood.

Here we describe a novel computational model of microvascular thrombus formation for detailed analysis of thrombus mechanics. Adopting a previously developed two-dimensional particle-based model focused on the thrombus shell formation, we revise it to introduce platelet agonists. Blood flow is simulated via computational fluid dynamics approach. In order to model soluble platelet activators, we apply Langevin dynamics to a large number of non-dimensional virtual particles. Taking advantage of the available data on platelet dense granule secretion kinetics, we model platelet degranulation as a stochastic agonist-dependent process.

The new model qualitatively reproduces enhanced thrombus formation due to granule secretion in line with *in vivo* findings and provides a mechanism for thrombin confinement at the early stages of aggregate formation. Our calculations also predict that release of dense granules results in additional mechanical stabilization of the inner layers of the thrombus. Distribution of the inter-platelet forces throughout the aggregate reveals multiple weak spots in the outer regions of thrombus, which are expected to result in mechanical disruptions at the later stages of thrombus formation.

## INTRODUCTION

Thrombus formation is an elaborate mechanism aimed at haemorrhage arrest that might otherwise cause excessive intratissular bleeding. However, unrestrained hemostatic response might lead to blood vessel occlusion and life-threatening embolism (Jackson, 2011). Platelets, being the principal structural elements of the arterial thrombus, rapidly respond to biochemical and biomechanical cues at the site of the injury and form the aggregate that is primarily stabilized by platelet-platelet interactions. Platelets largely define early mechanics of the thrombus outer shell (Babushkina et al., 2015, Kaneva et al., 2021), are vital for determining the occlusive scenarios (Ahmed et al., 2020, Belyaev et al., 2015) and contraction-driven re-arrangement of thrombus architecture (Nechipurenko et al., 2019), and provide a protected environment for blood coagulation to occur (Podoplelova et al., 2016, Tosenberger et al., 2013). Platelet activation is driven by receptor-mediated signaling, and soluble platelet agonists are now largely recognized as key regulators involved in arterial thrombus formation and stabilization (Sachs and Nieswandt, 2007).

For instance, thrombin and adenosine 5’-diphosphate (ADP) are potent inducers of platelet activation. Under the hemostatic response thrombin is produced in coagulation reaction cascade, whereas ADP is contained within platelet dense granules and is released through exocytosis along with multiple other compounds, including ATP, serotonin and polyphosphates. Experimental insights into heterogeneous organization of arterial thrombi gave rise to further understanding of the impact the platelet activators have on hemostasis (Stalker et al., 2013). Thrombin is believed to be almost entirely responsible for the formation of the thrombus core, the innermost layers of highly-activated platelets, whereas ADP and thromboxane A2 are principal regulators of the unstable thrombus shell assembly. How thrombin activity, platelet degranulation and agonist-induced platelet activation altogether orchestrate dynamics of thrombus formation is still unclear.

Theoretical modeling is widely used to study hemostasis due to the complexity of the system in question (Babushkina et al., 2015, Belyaev et al., 2015, Kaneva et al., 2021, Nechipurenko et al., 2019, Nechipurenko et al., 2020, Tosenberger et al., 2013, Xu et al., 2017, Yazdani et al., 2017). We have previously developed a particle-based model of arterial thrombus shell formation to address the state of adhesive receptors in different layers of the growing thrombi (Kaneva et al., 2021). However, since the model only considered the formation of thrombus shell, platelet activation was assumed to be an explicitly time-dependent process. To elaborate on the subject of thrombus heterogeneity, we modified the model and introduced explicit description of key platelet agonists. For the aim of this study, we adopted the platelet-platelet interaction mechanism and integrated it with the modeling approach we developed previously to describe platelet-flow interactions using quasi-steady flow approximation (Trifanov et al., 2018). The next step was supplementing this framework with a virtual particle model of agonist kinetics and a model of agonist-induced platelet activation.

Using the developed modeling framework, we demonstrate enhanced thrombus formation due to release of platelet dense granules. We also show that secretion provides additional mechanical stabilization of the inner layers of the thrombus. Using the model, we provide a possible explanation for the confinement of thrombin activity in space as being result of intrathrombus flows at the early stages of thrombus formation.

## MATERIALS AND METHODS

### Model framework

In the model, we consider the following processes that impact or drive thrombus formation: hydrodynamics of the blood flow, agonist-induced platelet activation, platelet-flow and platelet-platelet interactions, thrombin generation, secretion of ADP (through degranulation), and transport of thrombin and ADP (Fig. 1A).

**Figure 1.**
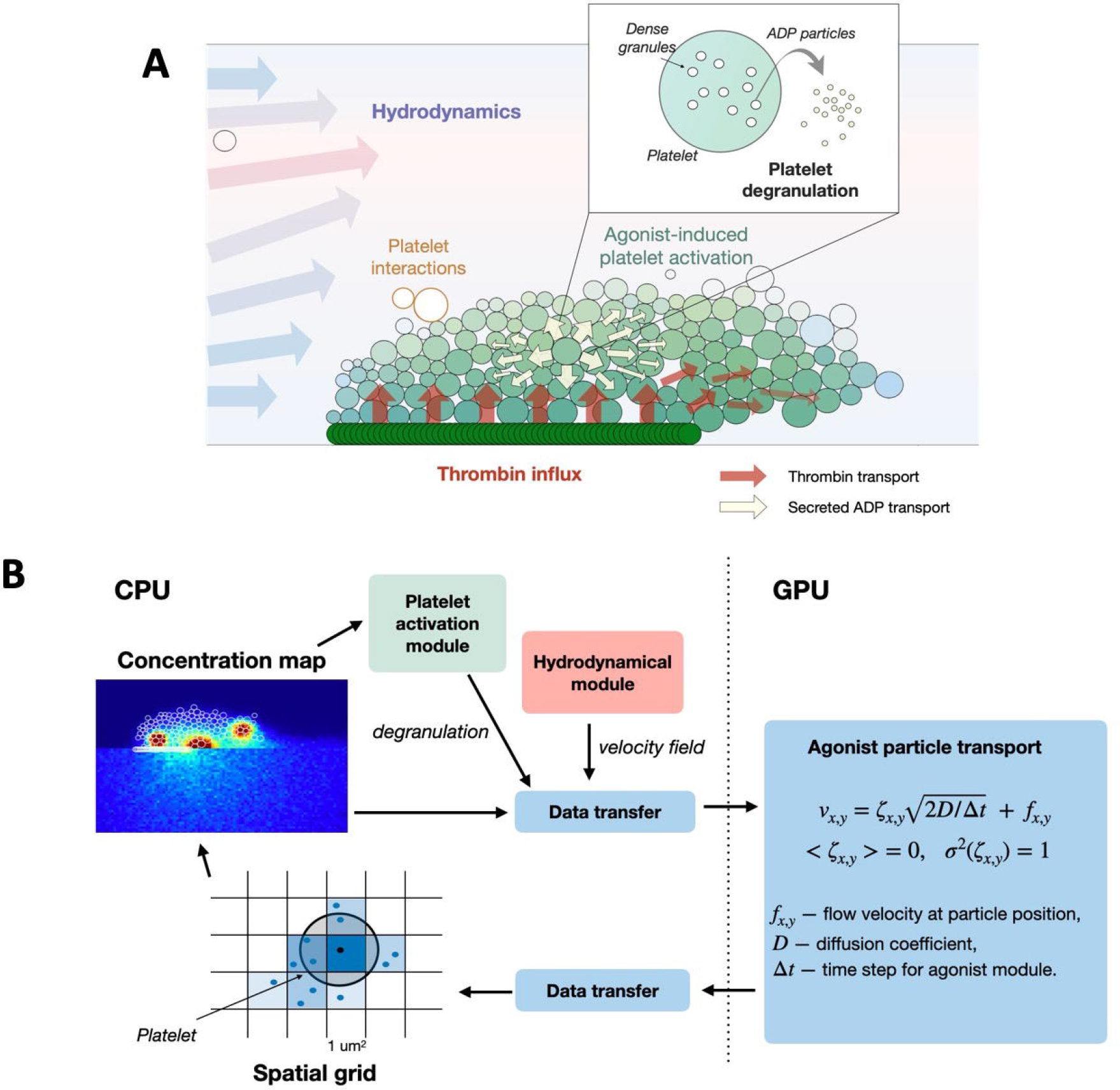
Model framework. (A) Diagram depicting interdependent modules within the model. The insert shows schematics of ADP generation through degranulation. (B) Data transfer between modules and separation of calculations into CPU- and GPU-hosted. Information about current positions of virtual particles together with velocity fields obtained from hydrodynamical module and degranulation signals is passed to GPU where positions of virtual particles are updated. Data is then transferred back to the CPU to calculate concentration map and update state of activation of platelets (see the text).

Hydrodynamics of the blood flow is described using quasi-steady flow approximation (see *Supplemental text A*) and blood (except platelets) is described is continuous media with Navier-Stokes equations. Platelets are described as particles traveling within the media, and platelets that constitute the aggregate are considered as stationary 2-dimentional discs with effective hydrodynamic radius. Agonists-induced platelet activation is described in a following manner: virtual particles that represent chemical species, change parameters of the mechanical interaction between platelets (see *Supplemental texts B* and *D* for details). Thrombin generation is described as a constant flux of thrombin-associated particles from the thrombus formation initiation region (the injury site). Secretion of ADP is described as generation of ADP-associated particles at the site of particular platelet. Transport of APD and thrombin is described using the Langevin dynamics of corresponding particles (see *Supplemental text C*).

The model comprises three distinct modules that interchange information about the current state of the system (Fig. 1B). Computational fluid dynamics (CFD) module recalculates pressure and velocity fields based on positions of platelets provided by particle dynamics (PD) module. Current hydraulic resistance in computational domain is used to update inlet parabolic flow boundary condition. Pressure and velocity fields obtained from CFD module by solving stationary Navier-Stokes equations are considered stationary in the PD module and used to calculate forces exerted on the platelets by the flow for the PD module. Information from CFD module is also used by agonist kinetics (AK) module to update concentration maps for agonists (see *Supplemental text E*). Platelet activation states are then updated in the PD module using concentration maps provided by AK module, and are in turn used to calculate inter-platelet forces. PD module is in the core of the model framework: here the positions of platelets are updated by numerical integration of the equations of motion, which comprise platelet-flow, platelet-wall and platelet-platelet forces and torques. Thrombin concentration also determines probability of degranulation for individual platelets, and signals for degranulation events are passed down to AK module to generate ADP.

The details on each module and simulation procedures are described in Supplemental texts A-E.

## RESULTS

### 1. Thrombin-driven hemostatic response leads to the formation of a stable platelet aggregate

To investigate the dynamics of thrombus formation driven solely by thrombin – a potent platelet agonist, we first analyzed the simplified case with a constant thrombin flux emanating from injury site as described in *Supplemental text C*. We also implemented explicit thrombin-dependent activation mechanism as described in *Supplemental text D* and simulated first 90 seconds of thrombus formation. Due to the stochastic nature of the model, this and following calculations were performed several times. Thrombus height was increased gradually, slowing down and reaching 12.25 ± 0.15 micrometers by the end of the simulation (Fig. 2A, black curve, Supplemental Video 1, 2). No thrombus disruptions were observed over the course of any simulation. The observed increase in stability of thrombus in comparison to disruptive thrombus shell (Kaneva et al., 2021) arises from the reinforcement of platelet-platelet bonds embedded in the model to reflect high potency of thrombin as platelet activator.

**Figure 2.**
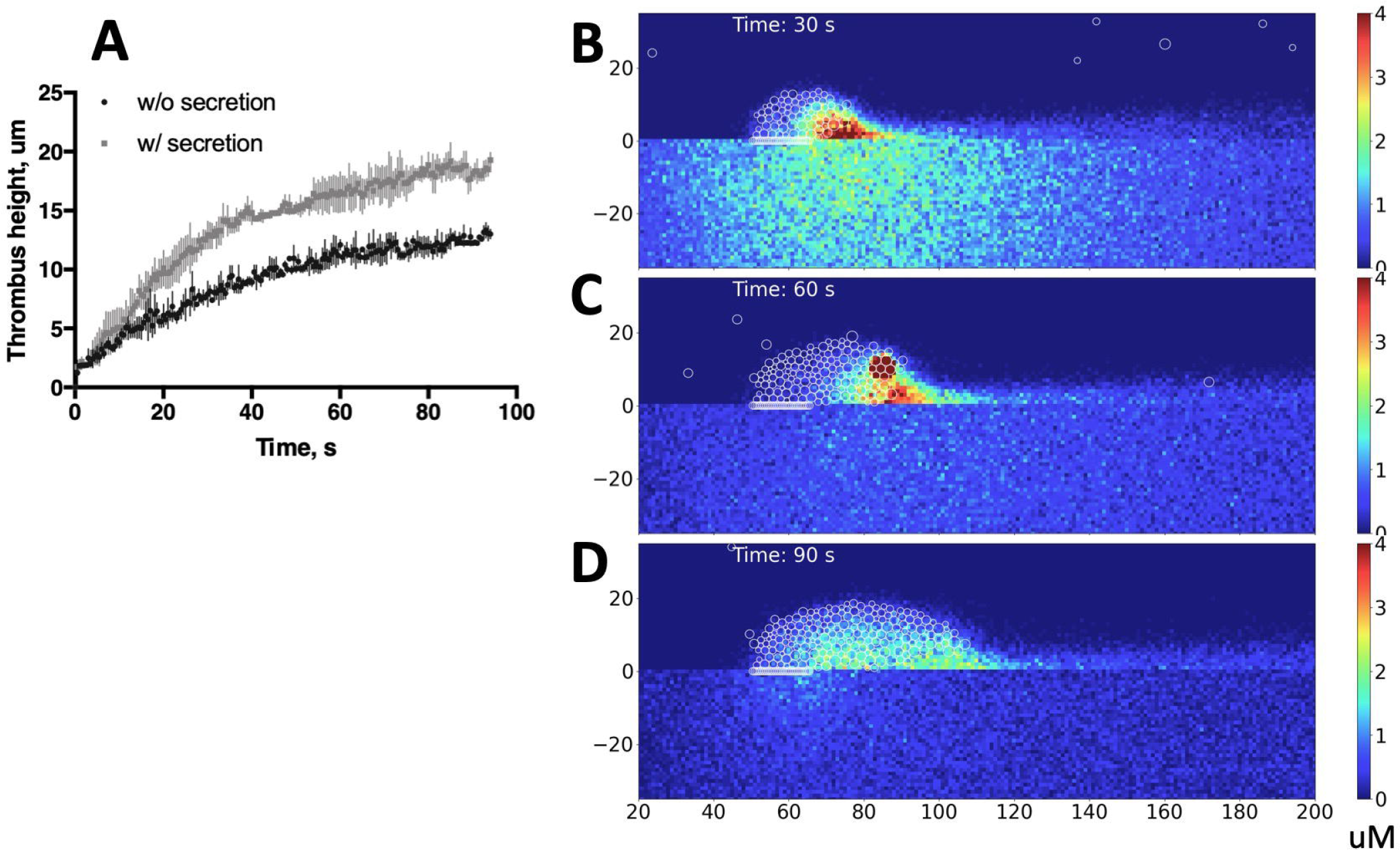
Dynamics of thrombus formation in the model. (A) Dynamics of thrombus growth as represented by thrombus height (in micrometers) versus model time (in seconds). Black curve corresponds to thrombin-only induced thrombus formation and grey curve is related to thrombus formation in the presence of both thrombin influx and platelet secretion. N=3 for each curve. (B-C) Snapshots of states of activation of platelets after 60 seconds of simulation for thrombin-induced thrombus formation (B) and thrombin and ADP-induced thrombus formation (C). First and second row show only the fraction of numerical value of the extent of activation induced by thrombin and ADP, correspondingly. Third row depicts overlapping of thrombin- and ADP-stimulated regions where platelets activated by both agonists are colored as a shade of green.

Thus, the model recapitulates formation of a stable thrombus “core” as observed in experiments, where only thrombin activity is retained.

### 2. Thrombin concentration gradient rapidly reaches steady state and limits thrombus propagation in the absence of granule secretion

We next examined whether spatial distribution of thrombin concentration is the limiting factor that determines the height of the thrombus in case when no other agonists are present. For this purpose, we analyzed the distribution of thrombin-particles in a large pre-formed platelet aggregate. Thrombin concentration decreases exponentially with distance from the injury site, and the gradient reaches steady state during first 3 seconds of simulation (Fig 3B), not much faster than does concentration gradient obtained in simulations with growing thrombus, described in a previous section (Fig. 3C). Comparing thrombin gradients from the two computational experiment settings, we conclude that it is thrombin gradient that limits the extent of thrombus growth in case when other agonists are absent. The mechanisms responsible for exponential decrease of thrombin concentration in space were further investigated and described below.

**Figure 3.**
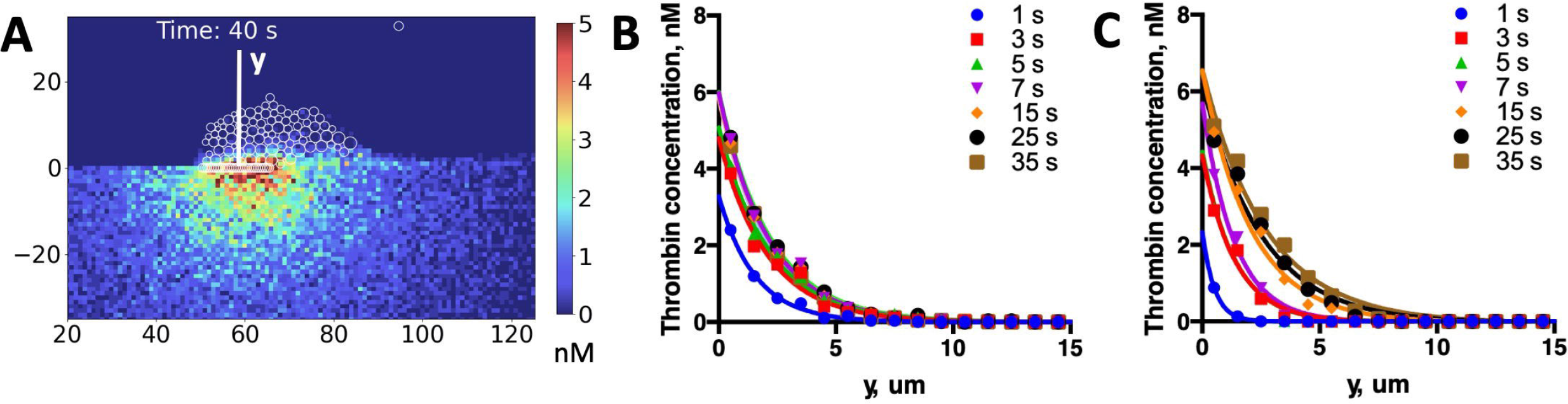
Thrombin concentration gradient. (A) Concentration map of thrombin in grown thrombus. Thrombin concentration gradient is shown for the center of the injury site denoted by white vertical axis *y* and its exponential fit as obtained for a preformed thrombus (B) and for a growing thrombus during its formation (C). Zero value at horizontal axis corresponds to the injury site. Different curves correspond to different stages of the simulation. In both representative simulations (B-C) concentration gradient rapidly reaches the steady state.

### 3. Platelet degranulation induced by thrombin enhances thrombus formation

Platelet dense granule secretion plays essential role in hemostasis as has been shown both *ex vivo* and *in vivo* (Israels et al., 1990, Meng et al., 2015). Disorders associated with abnormal degranulation result in impaired thrombus shell formation, such as in mice with Hermansky-Pudlak syndrome (Hermasky and Pudlak, 1969, Meng et al., 2015). To gain general understanding of the mechanism by which secreted agents affect formation of the thrombus, we focused on particular component of the dense granules – ADP and added platelet secretion to the model of thrombin-induced thrombus formation as described in *Supplemental text D*. Similar to the simulations described in the first section, we modeled 90 seconds of thrombus formation. Due to the burst-like character of granule secretion process (as separate granules are secreted), we observed burst-like kinetics of ADP generation and washout (Fig. 3B-D, Supplemental Video 4). Thrombus growth intensified compared to thrombin-only induced thrombus formation as seen by steeper increase in thrombus height and resulting thrombus height of 18.1 ± 0.4 micrometers compared to 12.25 ± 0.15 microns (Fig. 2A, grey and black curves). Considering that ADP gradient exceeds out of the thrombus region in the direction opposite the blood flow, we infer that this enhancement results from ADP-mediated platelet recruitment due to ADP-induced platelet activation, described in the *Supplemental text D*.

Thus, the described model demonstrates that release of ADP from platelet dense granules accelerates thrombus formation, in line with *in vivo* data.

### 4. High permeability of thrombus results in thrombin confinement effect due to convection-mediated dilution

To investigate the mechanisms responsible for thrombin confinement effect, we next analyzed the distribution and magnitude of plasma flows throughout the aggregate. In order to avoid non-realistic barriers for the flow due to the two-dimensional nature of the model, we decreased the hydrodynamic radius of each platelet during velocity field calculations (Fig. 4 A, C). The obtained geometry results in two-dimensional porosity of approximately 0.8, representing the upper bound of the 3d porosity measured for the outer layers of thrombus *in vivo* (Stalker et al., 2013). The heatmap of velocity magnitude values inside the aggregate at the 90 s of simulation demonstrates complex velocity field with an average value of 0.05 mm/s (Fig. 4 C, D). The estimate of the Peclet number for thrombin transport inside thrombus in current model exceeds unity, suggesting the significant effect of intrathrombus flow on thrombin concentration gradient, as evidenced in Figure 3A, B and thus explaining the exponential decrease of thrombin concentration as a function of distance from the injury site (Fig. 3 B, C). Importantly, the Peclet number for ADP transport is less than unity due to much higher diffusion coefficient and results in rapid diffusion-driven redistribution of ADP through the aggregate and hence – increased platelet recruitment at the outer regions of the aggregate.

**Figure 4.**
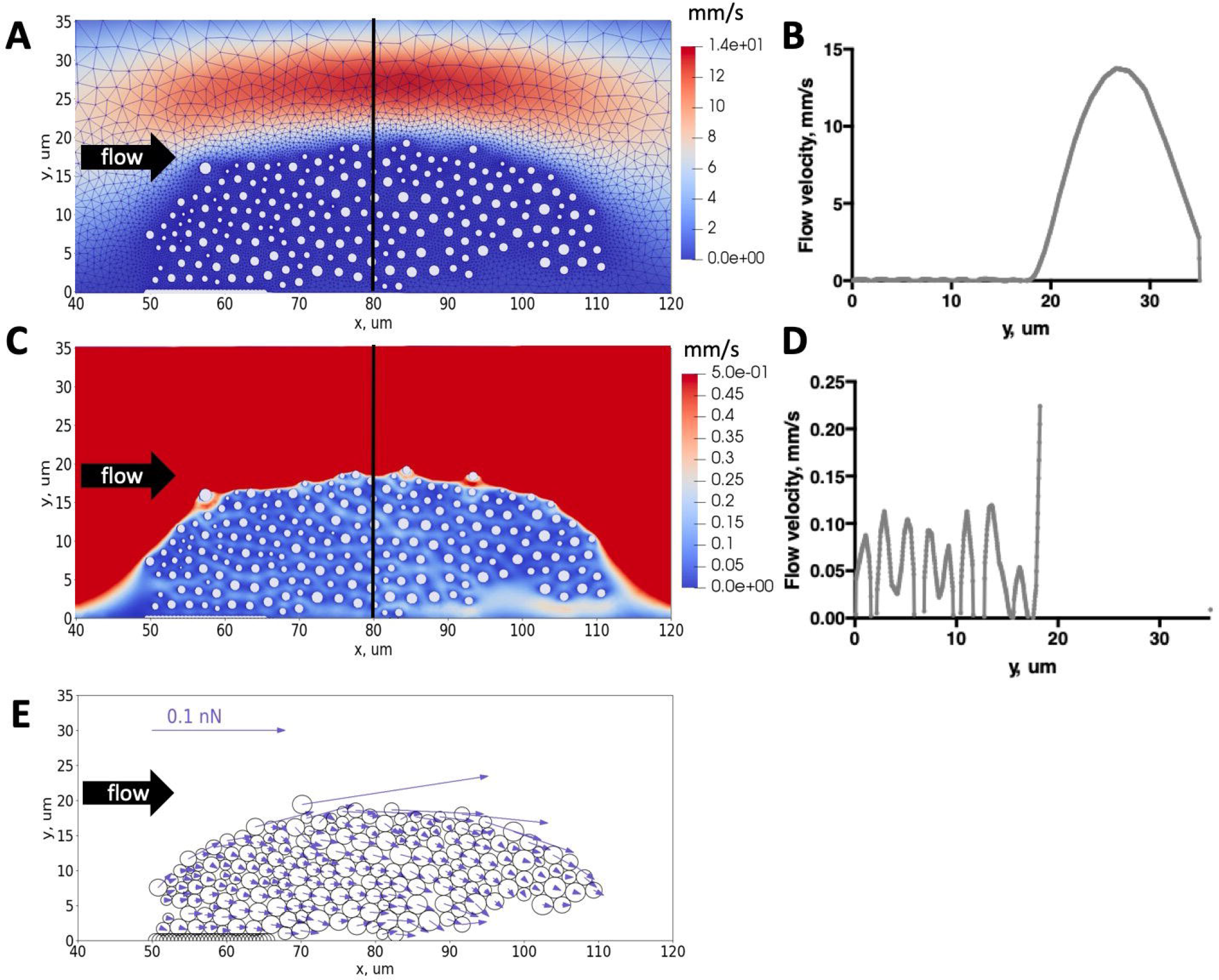
ADP concentration maps. (A-C) Spatial distribution of ADP at successive stages of platelet formation, at 30, 60 and 90 seconds of the simulation. Color corresponds to ADP concentration in uM. Degranulation events as one seen in (B) follow burst-like kinetics (see Supplemental Video 4).

### 5. Particle-based approach uncovers complex distribution of the tensile forces in heterogeneous thrombus

The complex dynamics of arterial thrombus observed *in vivo* is driven by the interplay between platelet-platelet and platelet-flow interactions. The values of critical tensile forces between platelets largely depend on the degrees of their integrins activation, which, in turn, depend on the local biochemical and biomechanical stimulus, experienced by these platelets.

To estimate the magnitudes and distribution of platelet-flow interactions we calculated the hydrodynamic drag forces within the aggregate. Analysis of the forces acting on the platelets in thrombus is depicted on Fig. 4E. Expectedly, these forces are much higher for the outer platelets of the aggregate, where the values of the shear rate are maximal.

To get deeper insight into the distribution of the mechanical forces, which stabilize the aggregate against the flow, we took advantage of the particle nature of the model and analyzed individual inter-platelet forces in thrombus. Interestingly, both repulsion and attraction forces between platelets show complex distribution with many “hot pairs”, wherein these forces are significantly increased (Fig. 5 B, C). The values of critical forces for given pair of platelets in our model is proportional to the product of platelet activation degrees (see *Supplemental text D*). The distribution of the critical tensile forces throughout the thrombus (Fig. 5A) qualitatively corresponds to the distribution of platelet activation levels (Supplemental Video 3). These forces are maximal at the base of the thrombus, where activation levels are higher, and decrease in the outer layers of the aggregate. To evaluate the safety margin S for each pair of platelets we used a simple formula:

**Figure 5.**
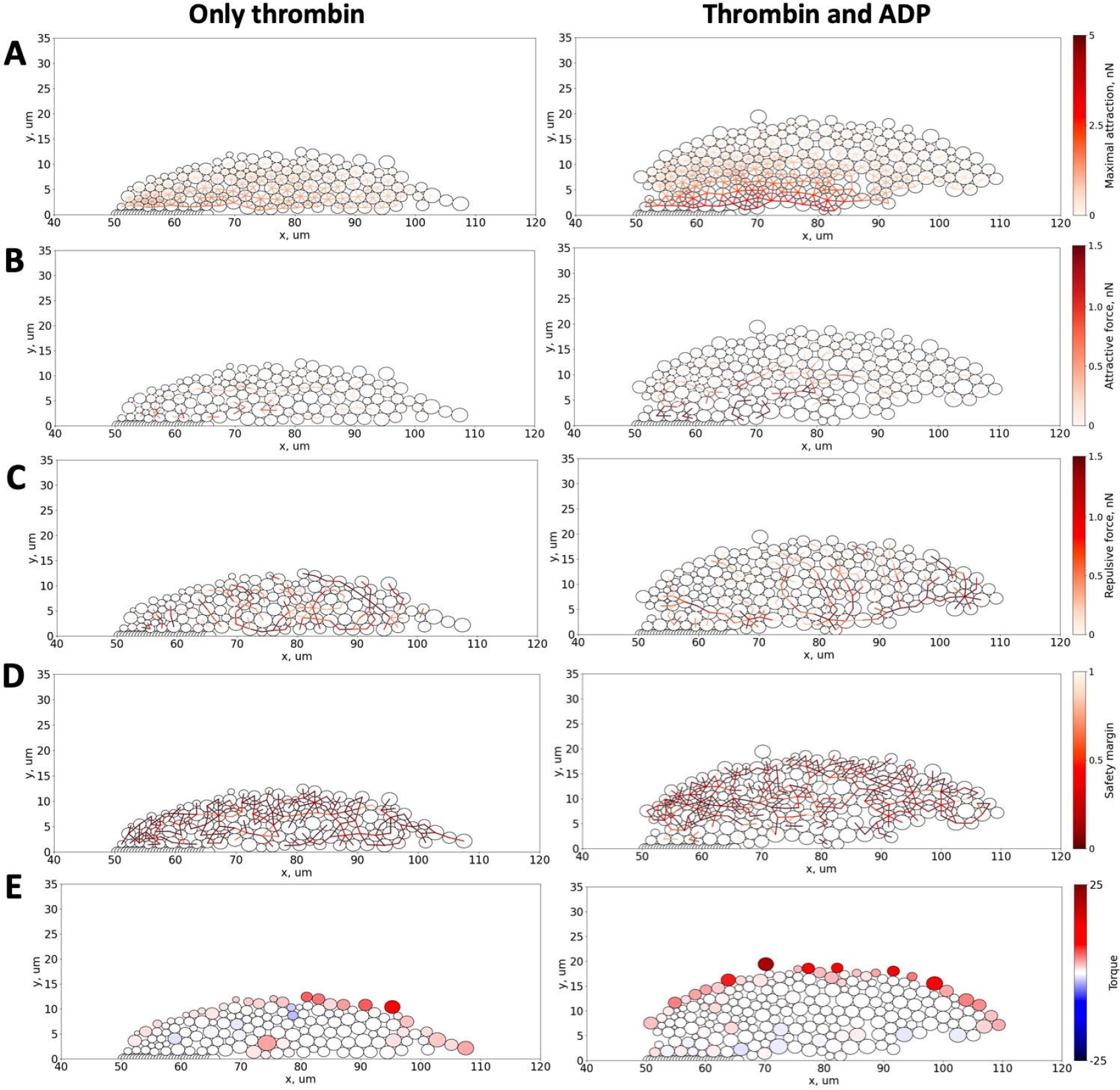
Intrathrombus plasma flows. (A) The velocity field and mesh in computational region. As thrombus propagates into the vessel, the flow becomes obstructed and inlet velocity decreases to maintain constant pressure drop in computational domain, though local flow velocity increases up to 14 mm/s compared to initial inlet velocity of 8.85 mm/s. This creates high shear rate at the boundary of the thrombus. Note that platelet radii are decreased (see the text). (C) Flow velocity profile taken in the center of aggregate denoted by black axis in (A). (B) Intrathrombus velocity field normalized to 0.5 mm/s and corresponding velocity profile. Velocity drops to zero inside platelets. (E) Hydrodynamic forces imposed on individual platelets. Platelets on the surface of the thrombus are exposed to the higher drag forces.

**Figure 6.** Distribution of inter-platelet forces through the aggregate. Left column corresponds to thrombus formed under thrombin-only stimulation, right column corresponds to thrombus formed under both thrombin and ADP stimulation. (A) Critical tensile forces between pairs of platelets as allowed for by their current state of activation. Platelets are centers of Morse potential field, and the force is proportional to the product of the extents of activation of interacting platelets. (B) Attraction forces between pairs of platelets which can be interpreted as tensile forces. (C) Repulsion between pairs of platelets. (D) Safety margin of a platelet-platelet bonds defined as S = 1 – F/F_max_, where F is attraction force between platelets and F_max_ is the critical force for this pair of platelets. Dark red bonds correspond to weak spots of thrombus where attraction forces are close to critical ones. (E) Torque imposed on individual platelets by their neighbors through stochastic springs representing vWF-mediated interaction. On the surface of the thrombus this torque counteracts the flow-induced torque and hence prevents lateral rolling of newly attached platelets under the shear stress.

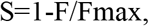

where F is the tensile force for given pair of platelets (derived from Morse potential), and F_max_ is the critical force for these platelets given their activation states.

Thus, the higher values of S (close to unity) correspond to the “safe” pairs – where current tensile forces are much lower than critical, while values closer to zero reflect “weak spots” – where platelet-platelet contact is likely to rupture in case if external forces are increased.

The distribution of the safety margin values throughout the aggregate (Fig. 5D), also demonstrates numerous “weak spots”. Interestingly, these weak spots are mainly located in the outer domain of the aggregate formed in the presence of both thrombin and ADP, but are still present in the innermost domain in case of thrombin-driven thrombus formation.

## DISCUSSION

Understanding the coupling between local biochemical/biomechanical stimuli and platelet-platelet interactions is crucial for elucidation of the basic mechanisms that regulate complex dynamics of arterial thrombus formation. Here we describe a novel *in silico* approach for investigation of the interplay between local biochemical stimuli, platelet-platelet interactions and the overall dynamics of thrombus under arterial blood flow.

In order to get deeper understanding of the processes that drive the formation of heterogeneous thrombus structure, we have significantly advanced the particle-based model, which was recently used to describe the plasticity of the arterial thrombus shell (Kaneva et al., 2021). The developed modeling framework considers the change of the blood flow due to formation of the thrombus, platelet integrin activation in response to thrombin and ADP, transport of thrombin and ADP, the secretion of ADP in response to thrombin, as well as two types of platelet-platelet interactions. The model was built in a bottom-up manner and its particle nature allows thorough and mechanistic analysis of the ongoing processes on the level of single platelets. However, such level of description results in high computational complexity of the model and thus makes it perfectly suitable for description of microvascular thrombosis on physiologically relevant timescales of several minutes.

Here we demonstrate that ADP, being secreted by platelets in response to thrombin, is rapidly distributed throughout the thrombus through diffusion process and leads to activation of the platelets that adhere to thrombus surface through primary stochastic interactions. Thus, dense granule secretion, which delivers ADP, results in significant increase of thrombus size, in line with *in vivo* findings (Meng et al., 2015).

Our results suggest a possible mechanism of thrombin confinement on the early stages of aggregate formation – when clot contraction is negligible. Intrathrombus plasma velocities result in advection-mediated dilution of thrombin, limiting the size of the thrombin-induced thrombus. Rapid stabilization of thrombin gradient inside thrombus is in line with experimental data obtained in mice: linear increase in thrombin sensor fluorescence observed over initial 120 s of thrombus growth *in vivo* suggests that total amount of thrombin in the thrombus remains constant over the indicated time period (Welsh et al., 2014).

Particle-based nature of the model allowed us to analyze the distribution of inter-platelet forces throughout the aggregate and conclude that dense granule secretion leads to additional stabilization of the innermost thrombus region. However, the ADP-dependent inter-platelet contacts in the outer layers of thrombus demonstrate multiple weak spots that might lead to thrombus disruptions at the later stages of thrombus formation.

The described model represents a unique computational instrument that couples local biochemical cues to the inter-platelet interactions and granule secretion. It exploits physically clear variables and parameters and is generally built in the bottom-up manner based on experimental data, including data on ADP-dependent activation of platelet integrins.

However, the current version of the model has several important limitations that will be addressed in future: it does not consider contraction of the thrombus, which is known to significantly reduce local aggregate permeability and hence alter the transport of species inside the aggregate (Tomaiuolo et al., 2014). The model does not explicitly account for coagulation reactions and fibrin formation, which might significantly impact local mechanical stability and thrombin transport. It does not consider such processes, as thromboxane A2 synthesis and release, and the activation of platelets by this substance (including granule release). We also did not account for additional platelet activation due to outside-in signaling (mechanically induced activation) or GPVI-induced signaling.

Despite these limitations, the developed instrument represents a valuable tool for the thorough analysis of the platelet mechanics that drives the dynamics of hemostatic response in arterioles and venules.

## ACKNOWLEDGMENTS

The work was supported by the Russian Foundation for Basic Research grant 19-51-15004 to F.A., by the Russian Science Foundation grant 20-45-01014 to M.A.P. and with the support of the RUDN University Strategic Academic Leadership Program for V.V. and by a grant from the endowment foundation “Science for Children”. The research was carried out using the equipment of the shared research facilities of HPC computing resources at Lomonosov Moscow State University. This research was performed within the framework of the Development Program of the Interdisciplinary Scientific and Educational School of Lomonosov Moscow State University «Photonic and Quantum technologies. Digital medicine». This project was also partly supported by the European Union’s Horizon 2020 research and innovation program (Grant No. 777072).

