## Supplemental Text for "Analysis of microvascular thrombus mechanobiology with a novel particle-based model"

**Supplemental texts.**

**Supplemental text A. Model geometry and hydrodynamics.**
As in our previous versions of particle-based models of thrombus formation [1, 2] we consider two-dimensional geometry to reduce model complexity. Blood vessel segment is modelled as a rectangular area with stiff boundaries. Initiation of thrombus formation takes place at a short section of the vessel wall covered by irreversibly adhered platelets (referred to as injury site below). Blood-borne platelets are considered as rigid discs which are capable of adhering to the injury site and to each other through a dual interaction mechanism (see *B. Platelet interactions*). Position and orientation of each platelet is fully defined by two spatial coordinates of the platelet center and the orientation angle. To account for experimentally observed platelet size heterogeneity [3], we impose Gaussian distribution to platelet radii as described in [1]. Soluble platelet agonists are represented by dimensionless virtual particles described by two spatial coordinates (see *C. Modelling platelet agonists*).
Blood plasma is modeled as a non-compressible Newtonian fluid described by Navier-Stokes and continuity equations. Employing method of computational fluid dynamics modelling described in [1], we consider a quasi-steady state approximation of the flow, updating boundary conditions and continuous pressure and velocity fields in case of significant thrombus shape change such as platelet attachment/detachment. To mimic availability of bypasses in hemodynamic network, we consider computational domain embedded in a linear segment of a hydraulic circuit such that their outlet ends coincide, and impose constant pressure drop condition on this segment. Since pressure drop *P* is proportional to resistance of hydraulic circuit element *R*, calculation of change in resistance of computational domain (due to thrombus formation) allows us to determine the maximum velocity value of the parabolic inflow ($v_0$):

$$R^i \sim \frac{P^i}{v_0^i},\quad P_t = \frac{P_{total} R^i}{R^i + bR_0}, \tag{1,2}$$

$$v_0{}^{i+1} = \frac{v_0{}^i P_t}{P^i}, \tag{3}$$

where $R^i$ and $P^i$ are resistance and pressure drop in the computational domain on the $i^{\text{th}}$ iteration, $P_t$ is the targeted pressure drop in the computational domain, $P_{total}$ is the constant pressure drop within prolonged circuit segment, $b$ is the ratio of the difference in lengths of circuit segment ($L_{dP}$) and computational domain (X) to the computational domain length (X), and $R_0$ is the initial resistance of the vessel without the thrombus (computational domain). $v_0{}^{i+1}$ is passed to CFD module to obtain new velocity and pressure fields in the computational domain, and the iteration process stops when difference between obtained and targeted pressure is less than 1 percent of the targeted pressure.
Vessel walls and surfaces of platelets within thrombus are no-slip boundaries. These platelets are considered as impermeable flow barriers; to allow permeability of the thrombus, hydrodynamical radii of these platelets are two times smaller than geometrical ones (which are used to calculate interplatelet forces). Flow disturbances created by free-flowing platelets are neglected: thus, the current model employs one-way coupling approximation for moving platelets, as they experience Stokes force from the fluid, but don't affect the velocity field. For simplicity, we assume that average flux of the incoming platelets at the inlet boundary is spatially uniform. However, parabolic flow profile gives rise to platelet margination effect reported in experiment [4]. Platelet concentration was determined as described in [2].

**Supplemental text B. Platelet interactions.**
In the model there are several forces exerted on the platelets, arising from platelet-wall, platelet-flow and platelet-platelet interactions (see [2] for details). Since acquirement of shell instability requires a dual interplatelet interaction mechanism, as has been shown in [2], we adopt it without modifications. Such mechanism recognizes primary glycoprotein-Ib-mediated and secondary

integrin-αIIbβ3-mediated platelet tethering, the former described by stochastically associating and dissociating springs and the latter by a deterministic Morse potential.
Reflecting the distinctive ability of platelets to gradually respond to stimuli, the stability of platelet association is assumed to be regulated by local agonist concentrations. We interpret this property as an extent of platelet activation and assign a corresponding numerical parameter α to each platelet. The exact manner of its determination is described in *D. Agonist-induced platelet activation*.

### Supplemental text C. Modelling platelet agonists.

In order to model soluble platelet activators with nearly single-molecule precision we apply Langevin dynamics to a large number of non-dimensional non-interacting massless virtual particles which represent small amounts of substance (Fig. 1B); this approach allows us to spatially and temporally resolve agonist transport. To account for possibility of substance deposition in the surrounding tissue, we consider two domains of virtual particles transport: the vessel domain that coincides with the hydrodynamics computational domain and the subendothelium domain. Once generated, a particle is subject to diffusion and advection in the vessel region and only diffusion in the subendothelium region; particles that escape the transportation domains are eliminated (see *E. Numerical calculations*). Since the model is two-dimensional and does not allow transportation freedom reserved by three-dimensional packing of spherical objects, platelets are assumed to be permeable for virtual particles, which is achieved by imposing null flow velocity field condition on intraplatelet regions. Local activator concentration that affects a specific platelet is proportional to total number of virtual particles in a 1-µm grid element that corresponds to the center of the platelet. Further refinement of the mesh would result in an unjustified assignment of local agonist concentration to a platelet which is two micrometers in diameter on average. Particles corresponding to thrombin and ADP follow different rules of generation and degradation.

Thrombin production culminates a complex set of coagulation reactions that take place on cell membranes as well as in the media. As here we do not focus on kinetics of thrombin production, we neglect the coagulation cascade and assume that thrombin enters the vessel in a constant influx of 0.1 nmol per $m^2$ per s. Moreover, increase in prothrombin concentration in blood plasma does not lead to enhanced arterial thrombus growth [5]. Since the income of direct thrombin precursor has no apparent effect on thrombin generation, we suggest that this process is confined to the area adjacent to the injury site and is rate-limited (i.e. thrombin flux is determined by the number of prothrombinase molecules at the injury site, but not the substrate transport). To account for thrombin inhibition by plasma-borne inhibitors, we assume that thrombin particles are randomly degraded at rate of 0.02 particle per second [6].

ADP particles only originate from platelet dense granule upon their release (see Db. *Thrombin-induced platelet degranulation*). Unlike thrombin particles, these particles are not extinguished due to hydrolysis since the adenine nucleotide catabolism reactions time scale in human blood [7] is much larger than characteristic time of ADP washout in the considered geometry scale.

### Supplemental text D. Agonist-induced platelet activation.

#### a. Effect of agonists on platelet adhesion and aggregation.

Understanding coupling between adhesive properties of platelet and its state of cellular activation is of pivotal importance for describing mechanical aspects of platelet aggregation and thus thrombus formation. We adopt a universal variable α, namely extent of activation, to account for both interplatelet interaction mechanisms. The value of α in the model can be interpreted as the degree of platelet integrin activation. The maximum integrin-dependent interplatelet forces (the critical force) between two platelets is derived from the Morse potential and is proportional to the product of their activation extents.

Since intrinsic activatory pathways are triggered by soluble agonists, we assume that this property has to depend on local agonist concentration. Moreover, due to irreversibility of some activation-associated events, we have to consider that current level of platelet activation depends on the preceding exposure to agonists.

Physiological activation of platelets is differentially regulated by thrombin and ADP; therefore, we assume the same for their adhesive properties and model thrombin-induced and ADP-induced extent of activation according to specific characteristics of these processes. Thrombin is known to be a potent platelet agonist that causes rapid irreversible activation of platelets. For simplicity, we assume thrombin-induced activation to be irreducible and explicitly concentration-dependent. We assume dependence of extent of activation parameter on thrombin concentration to resemble Hill equation for dose-dependency for non-cooperative conditions. Constant of half-response is arbitrary chosen to fall within physiological range. We introduce a delay time to account for time needed for signal transduction pathways to activate platelet integrins after exposure to thrombin. This delay is mimicked by a continuous dependence of the extent of activation on the time since initiation of the activation. Once achieved extent of activation is not reducible.

$$\alpha_{\text{IIa}}(t+\Delta t)=\max\left\{\frac{\alpha_{\text{IIa,max}}}{1+\frac{[\text{IIa}]_{\frac{1}{2}}}{[\text{IIa}]}}\frac{1}{1+\frac{\tau}{t_{in}}},\ \alpha_{\text{IIa}}(t)\right\}, \tag{4}$$

where $t_{in}$ — time spent in thrombus, [IIa] — thrombin concentration, $\alpha_{max}$ — maximal extent of activation, $[\text{IIa}]_{1/2}$ — half response concentration, $\tau$ — characteristic time of activation onset.

The maximal extent for this type of activation (thrombin-mediated activation) exceeds the level corresponding to 100% integrin activation in order to account for additional stabilizing effect caused by thrombin-induced fibrin net formation. In case when 2 maximally activated platelets (by thrombin) interact with each other, their critical force would be 5 nN.

To model reversible ADP-induced activation we consider fibrinogen binding kinetics in response to ADP observed in the experiment [8] and determine change in the number of occupied fibrinogen receptors, assuming that number of receptors available for binding, i.e. activated integrins, depends on local ADP concentration. We then define extent of activation as fraction of fibrinogen-bound integrins for a specific platelet: $\alpha_{ADP} = [R{:}Fb]/R_{total}$, where $[R{:}Fb]$ is number of occupied integrins and $R_{total}$ is total number of integrins per platelet [9], and solve a differential equation derived from fibrinogen binding kinetics to obtain its current value. The main assumptions of this consideration are as follows:

1. Fibrinogen binding to platelet integrins in response to ADP occurs in accordance with the law of mass action in the excess of the ligand (Fb). Integrins available for binding are denoted as R. Values of the rate constants are obtained from [8], fibrinogen concentration in blood $[\text{Fb}]_0$ is constant [10]:

$$Fb + R \rightleftharpoons R{:}Fb, \tag{5}$$

$$\frac{d[R{:}Fb]}{dt} = k_{on}[Fb]_0[R] - k_{off}[R{:}Fb]. \tag{6}$$

2. Since extent of activation has to depend on ADP concentration, and number of integrins available for binding [R] is the only variable parameter in Equation 6, we assume that at each given time it depends on ADP concentration. To determine nature of this relation, we refer to ADP concentration dependence curve for fibrinogen binding [8] and fit it with sigmoidal function (Supplemental Figure 1). This yields an expression for number of activated integrins (available for binding and already occupied, [R] + [Fb:R]) as a fraction of maximal possible number of integrins to be activated by ADP $R_{max}$. To respect concurrent thrombin-induced fibrinogen binding, we

interpret extent of activation induced by thrombin as an additional fraction of already occupied integrins, therefore inaccessible for ADP-induced binding. Corresponding number of integrins is then subtracted from maximal possible number of integrins to be activated by ADP:

$$[R] = \frac{R_{max} - \alpha_{IIa} R_{total}}{1 + \frac{[ADP]_{1/2}}{[ADP]}} - [R:Fb], \tag{7}$$

where $\alpha_{IIa}$ – thrombin-induced extent of activation of a platelet, $[ADP]_{1/2}$ — characteristic ADP concentration. Thus, resulting expression for $\alpha_{ADP}$ is as follows:

$$\alpha_{ADP}(t + \Delta t) = \alpha_{ADP}(t) + \Delta t \left[ k_{on}[Fb]_0 \left( \frac{\frac{R_{max}}{R_{total}} - \alpha_{IIa}(t)}{1 + \frac{[ADP]_{\frac{1}{2}}}{[ADP](t)}} \right) - k_{off}\alpha_{ADP}(t) \right]. \tag{8}$$

3. A ligand-bound integrin is stabilized in an activated state, i.e. decrease in ADP concentration which might lead to inactivation of this integrin will not affect it until it is occupied by fibrinogen; upon fibrinogen dissociation it changes its state to closed instantly. Here we suggest that slow fibrinogen dissociation rather than intracellular signaling is the rate-limiting process.

ADP- and thrombin-induced extents of activation sum to total extent of activation used to determine mechanical properties of interplatelet interactions:

$$a = \alpha_{IIa} + \alpha_{ADP}. \tag{9}$$

### b. Thrombin-induced platelet degranulation.

Aiming at accurate characterization of ADP generation in the model we took advantage of the available data on dense granule secretion kinetics [11], specifically time course of thrombin-induced secretion in suspension of washed platelets as measured through ATP release. First, we hypothesized that release of a single granule is a random event and therefore provided constant thrombin activity secretion is a pure-death process of granule pool elimination and can be described by two parameters, probability density for release of a single granule $p$ and fraction of granules to be potentially released $S_{max}$:

$$\frac{N_{DG}(t)}{N_{DG}(t = 0)} = S_{max}(1 - e^{-pt}). \tag{10}$$

We were able to fit experimental data on secretion time course (N=5) within the margin of the error (Supplemental Figure 2). For further reasoning we assume that $S_{max}$ does not depend on thrombin concentration, which is consistent with experimental data on secretion kinetics in case of ten-fold increase in thrombin activity: overall fraction of released granules does not increase while the rate of the process is affected significantly [12]. To determine the nature of dependency of $p$ on thrombin activity $[IIa]$, we approximated dose-dependency secretion curves (N=5, at time $t_0$ = 1 min) by a sigmoidal function with parameter $A$ (Supplemental Figure 3):

$$\frac{N_{DG}(t = t_0)}{N_{DG}(t = 0)} = S_{max}(1 - e^{-pt_0}) = S_{max} \frac{[IIa]^2}{A^2 + [IIa]^2}. \tag{11}$$

Solving this equation for $p = p([IIa])$ we get explicit expression:

$$p([IIa]) = \frac{1}{t_0}\left(1 + \frac{[IIa]^2}{A^2}\right). \tag{12}$$

To convert thrombin concentration in nM calculated in agonist kinetics module to thrombin activity in U/ml used in degranulation module we impose a factor of 0.1 on the value of thrombin concentration. Number of granules per platelet is proportional to platelet volume, which is a heterogeneous property itself. Mean number of granules per platelet equals 19 reflecting recent findings for murine platelets [13]. Platelets release their granules independently to the extent of $S_{max}$.

Thus we introduce platelet degranulation into the model as a stochastic single-granule scale process, probability density function of which depends non-linearly on thrombin concentration.

**Supplemental text E. Numerical calculations.**

CFD calculations are performed using simpleFoam solver provided by OpenFOAM software [14]. Recalculation of velocity and pressure fields are triggered by attachment/detachment of platelets or their relocation within thrombus that exceeds 1 micrometer, but CFD update is not initialized more often than once in 2 ms model time. Particle dynamics recalculation is run in parallel using OpenMP. Equations of motion are solved using modified version of Verlet algorithm [15].
AK module converts the fine-mesh solution for velocity field provided by CFD module to a coarse grid to accelerate determination of velocities corresponding to positions of virtual particles present in large numbers. To satisfy high computational resources demand of virtual particles dynamics calculations, update of particle positions is implemented on CUDA-enabled GPUs. Information about current position of all virtual particles and corresponding velocities is transferred to GPU, where stochastic members of Langevin equation are calculated and equations of motions themselves are solved. Yielded positions for complete set of virtual particles are transferred back to CPU to be converted to concentration maps as described above and fed to PD module. Timestep for AK module is 0.2 ms model time while timestep for PD module is 0.0002 ms (see Supplemental Table for other model parameters). Single ADP particle represents 90 ADP molecules and, considered in a 1x1x1 micrometer grid element, corresponds to approximately 150 nM of substance concentration, which determines minimum resolved change in ADP concentration. Note that despite we generally consider model as 2d, in order to get the estimates of 3d concentrations and forces acting on single platelets, we consider our 2d domain as a 1 micrometer-thickness slice of an effective 3d system. The depth of 1 micrometer corresponds to the typical platelet size.

Assignment of 1 thrombin virtual particle to 1 thrombin molecule would result in minimum resolved change in thrombin concentration of 1.67 nM, an amount capable of invoking significant response from platelets. Thus, we attribute four thrombin virtual particles to a single thrombin molecule, reducing minimum resolved amount of thrombin to 0.42 nM.
Elimination of virtual agonist particles upon their escape out of the subendothelium domain leads to underestimation of the concentration and is a limitation of the current model. To minimize this effect, we consider subendothelium domain to be twice as large as the vessel domain, since increase in the size of buffering zone reduces the resulting error.
Program code unrelated to CUDA-hosted computation was written in C++. Calculations were performed using Lomonosov-2 supercomputer resources at Lomonosov Moscow State University.

**Supplemental Table 1. Model parameter values.**

| Parameter | Symbol | Value | Units | Reference |
|---|---|---|---|---|
| Time step for particle dynamics module | dt | 0.002 | ms | This model |
| Time step for agonist kinetics module | | 0.2 | ms | This model |
| Thrombin virtual particles per molecule | | 4 | | This model |
| ADP virtual particles per molecule | | 0.011 | | This model |
| ADP virtual particles per dense granule | | 10500 | | This model |
| Mean number of dense granule per platelet | $N_{DG}$ | 19 | | [12] |
| Diffusion coefficient for thrombin | $D_{IIa}$ | 67 | $um^2/s$ | |
| Diffusion coefficient for ADP | $D_{ADP}$ | 257 | $um^2/s$ | |
| Geometry and hydrodynamics | | | | |
| Vessel height (height of computational domain) | H | 35 | μm | [2] |
| Length of computational domain | X | 150 | μm | |
| Dynamic viscosity | η | $10^{-3}$ | Pa * s | |
| Dynamic viscosity for rotation | $\eta_w$ | $10^{-15}$ | N * s | |
| Near wall shear rate | $\gamma_0$ | 1000 | $s^{-1}$ | This model |
| Injury length | L | 15 | μm | This model |
| Pressure drop | $P_{total}$ | 50 | Pa | This model |
| Length of constant pressure drop domain | $L_{dP}$ | 750 | μm | This model |
| Platelets and platelet interactions | | | | |
| Platelet mass | m | $4 * 10^{-15}$ | kg | [2] |
| Platelet radius | R | 1 | μm | |
| 2D platelet concentration (number per computational domain) | $n_0$ | 5 | | |
| The rigidity of platelet interaction with the vessel wall | $k_{wall}$ | 1.5 | nN/μm | |
| The rigidity of platelet-platelet interaction | $k_{el.repulsion}$ | 0.5 | nN/μm | |
| Stochastic spring constant | $k_{ST}$ | 30 | pN/μm | |
| Spring equilibrium lenght | $L_0$ | 0.05 | μm | |

| | | | | |
|---|---|---|---|---|
| Constant, characterizing the dependence of the stochastic spring creation probability on the distance between platelets | $c_{as}$ | 10 | $\mu m^{-2}$ | |
| Maximal rate of bond creation | $k_0$ | 9 | $ms^{-1}$ | |
| Ratio of minimal to maximal probability of bond creation | $\delta^2$ | 0.001 | | |
| Rate of bond dissociation with no force applied | $k_{0,dis}$ | 17.1 | $s^{-1}$ | |
| Constant, characterizing the dependence of the bond break probability on the applied force | $\beta$ | 1.358 | $nN^{-1}$ | |
| Constant, characterizing the dependence of integrins-mediated force on the distance between platelets | $\lambda_M$ | 20 | $\mu m^{-1}$ | |
| Constant, characterizing the maximum integrins-mediated force between activated platelets | $A_M$ | 10 | nN | |
| ADP-induced aggregation | | | | |
| ADP concentration resulting in half integrin availability | $[ADP]_{1/2}$ | $1.5\ 10^{-3}$ | mM | [7] |
| Maximal number of integrins ADP can make available for binding | $R_{max}$ | 60000 | | |
| Fibrinogen binding apparent $k_{on}$ | $k_{on}(Fb)$ | $141.5\ 10^{-6}$ | $mM^{-1}\ s^{-1}$ | |
| Fibrinogen binding apparent $k_{off}$ | $k_{off}(Fb)$ | 0.018 | $s^{-1}$ | |
| Fibrinogen concentration | $[Fb]_0$ | 2 | g/l | [8] |
| Total amount of integrins | $R_{total}$ | 80000 | | [9] |
| Thrombin-induced aggregation | | | | |
| Thrombin concentration resulting in half platelet activation | $[IIa]_{1/2}$ | 5 | nM | This model |
| Thrombin activation time-dependence parameter | $\tau$ | 2 | s | This model |
| Thrombin inflow | | | | |
| Thrombin influx at the injury site | | $10^{-20}$ | $mM\ um^{-2}\ s^{-1}$ | This model |
| Dense granule release | | | | |
| Probability density expression parameter | A | 0.95 | nM | Estimated on the basis of experimental data [10] |
| Maximum fraction of granules to be released | $S_{max}$ | 0.87 | | Estimated on the basis of experimental data [10] |

**Legends for supplemental figures and videos**

**Supplemental Figure 1. Characterization of fibrinogen binding in response to different ADP concentrations.** Dots correspond to experimental data reported in [8]. See Supplemental text Da.

**Supplemental Figure 2. Time-course of dense granule secretion in response to thrombin.** Dots correspond to the data reported in [11] and represent mean ± SD for 5 experiments using wild-type mice. See Supplemental text Db.

**Supplemental Figure 3. Effect of thrombin activity on dense granule secretion.** Dots correspond to the data reported in [11] and represent mean ± SD for 5 experiments using wild-type mice. See Supplemental text Db.

**Supplemental Video 1. Thrombin-induced thrombus formation.** Platelets are shown as color-filled circles (the higher the extent of activation, the darker the blue). Virtual particles that represent the injury site are shown as dark green circles.

**Supplemental Video 2. Evolution of thrombin concentration map during thrombin-induced thrombus formation.** Platelets and virtual particles that represent the injury site are shown as white circles. Vessel domain resides in the upper part of computational domain. Spatial distribution of thrombin concentration is converted into a heatmap, with blue denoting absence of thrombin and red denoting concentration higher than 5 nM. Legend values are given in nM. Borderline between vessel domain and subendothelium domain is not shown.

**Supplemental Video 3. Thrombin and ADP-induced thrombus formation.** Platelets are shown as color-filled circles (the higher the extent of activation, the darker the color). Blue coloring corresponds to thrombin-induced activation, yellow coloring corresponds to ADP-induced activation, overlay of two types of activation results in green coloring. Virtual particles that represent the injury site are shown as dark green circles.

**Supplemental Video 4. Evolution of ADP concentration map during thrombus formation.** Platelets and virtual particles that represent the injury site are shown as white circles. Vessel domain resides in the upper part of computational domain. Spatial distribution of ADP concentration is converted into a heatmap, with blue denoting absence of ADP and red denoting concentration higher than 4 uM. Legend values are given in uM. Borderline between vessel domain and subendothelium domain is not shown.
