## Supplementary figures and images for "Analysis of microvascular thrombus mechanobiology with a novel particle-based model"

### Supplemental Figure 1

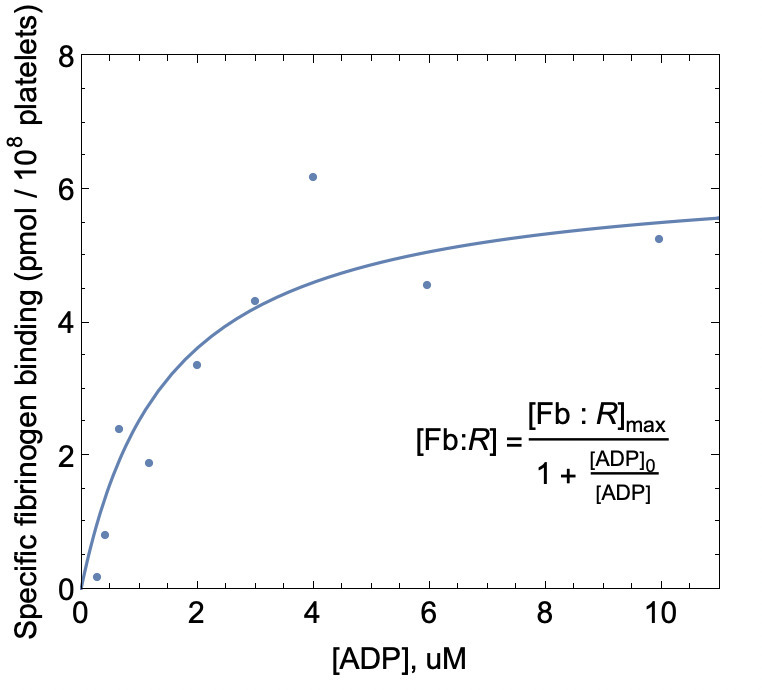

### Supplemental Figure 2

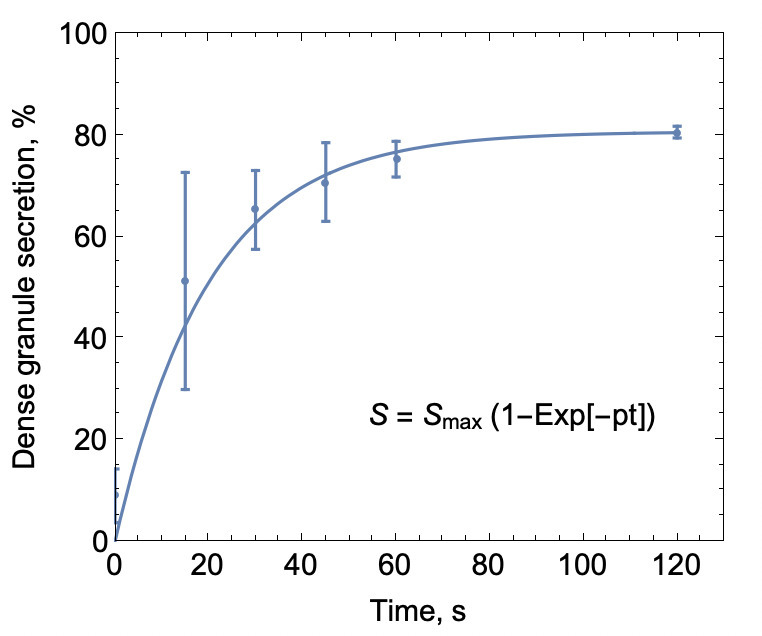

### Supplemental Figure 3

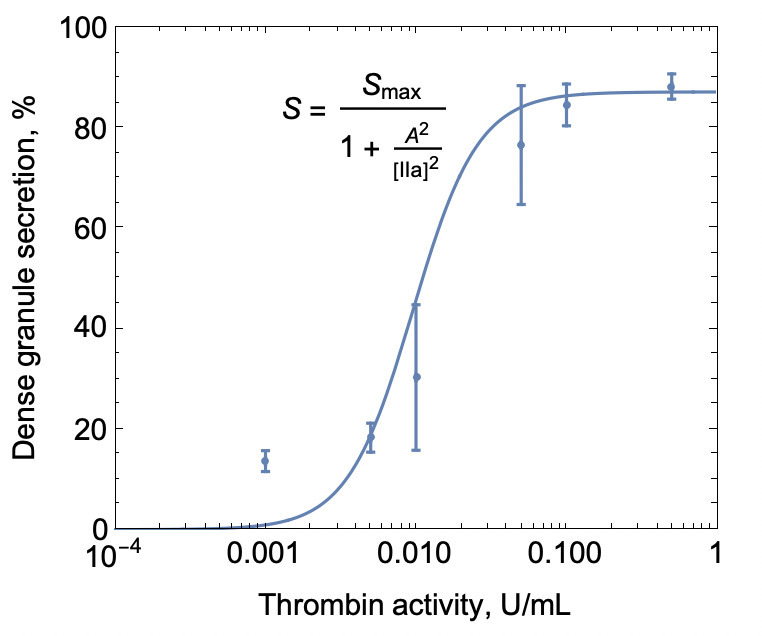
